# Diffusion Tensor Tractography of Brainstem Fibers and Its Application in Pain

**DOI:** 10.1101/569723

**Authors:** Yu Zhang, Andrei A. Vakhtin, Jennifer S. Jennings, Payam Massaband, Max Wintermark, Patricia L. Craig, J. Wesson Ashford, David J. Clark, Ansgar J. Furst

## Abstract

Evaluation of brainstem pathways with diffusion tensor imaging (DTI) and tractography may provide insights into pathophysiologies associated with dysfunction of key brainstem circuits. However, identification of these tracts has been elusive, with relatively few in vivo human studies to date. In this paper we proposed an automated approach for reconstructing nine brainstem fiber trajectories of pathways that might be involved in pain modulation. We first performed native-space manual tractography of these fibers in a small normative cohort of participants and confirmed the anatomical precision of the results using existing anatomical literature. Second, region-of-interest pairs were manually defined at each extracted fiber’s termini and nonlinearly warped to a standard anatomical brain template to create an atlas of the region-of-interest pairs. The resulting atlas was then transformed non-linearly into the native space of 17 veteran patients’ brains for automated brainstem tractography. Lastly, we assessed the relationships between the integrity levels of the obtained fiber bundles and pain severity levels. Fractional anisotropy (FA) measures derived using automated tractography reflected the respective tracts’ FA levels obtained via manual tractography. A significant inverse relationship between FA and pain levels was detected within the automatically derived dorsal and medial longitudinal fasciculi of the brainstem. This study demonstrates the utility of DTI in exploring brainstem circuitries involved in pain processing. In this context, the described automated approach is a viable alternative to the time-consuming manual tractography. The physiological and functional relevance of the measures derived from automated tractography is evidenced by their relationships with individual pain severities.

## Introduction

The brainstem, including the midbrain, pons, and medulla, involves structures with complex white matter pathways and gray matter nuclei that are concentrated in a small area. Intricate brainstem circuitries and nuclei serve systems such as respiratory and cardiovascular regulation, sleep and alertness, pain, posture, mood, and mnemonic functions. It is crucial to understand how structural changes in small brainstem regions and circuitries may cause/alter pathologies.

Studies of brainstem substructures using anatomical brain MRI have been complicated by difficulties in detecting neuronal loss due to lack of sufficient contrast to delineate small internal substructures in intensity-based images. Diffusion tensor imaging (DTI) is a noninvasive MRI imaging technique that measures changes of water diffusion in white matter microstructures. Fractional anisotropy (FA), one of the standard DTI indices, is known to be sensitive to detect damages in orientationally organized structures (e.g. white matter fibers). Furthermore, based on computing the directional information in each voxel of DTI, tractography is used to reconstruct trajectories of white matter tracts that correspond to known neuroanatomy in 3-dimensional space [1,2]. Diffusion tensor tractography has orientation-based contrasts and thus permits not only the anatomical illustration of neural pathways, but also examines the integrity and structural connectivity of a specific pathway by measuring the microstructural or fiber indices along the reconstructed tracts.

Diffusion tensor tractography has been broadly applied in neurosurgical settings, such as navigating tumor resections to avoid damage to surrounding vital neural pathways [3] and providing guidance to targets of deep brain stimulation electrodes [4]. Quantitative DTI measures in tractography-derived fiber bundles have also been used for detecting microstructural deficits in multiple neurologic and psychiatric disorders, such as amyotrophic lateral sclerosis [5], cognitive impairments and Alzheimer’s Disease [6–8], and other neurologic and psychiatric disorders [9]. However, tracking and isolating brainstem pathways using DTI is challenging due to their spatial overlapping and crossing with other major tracts that project to cerebral cortices. Previous tractography investigations pertaining to the brainstem have focused on major fiber bundles between the brainstem and the cortex [10] that provided few details about specific fiber tracts within the brainstem. A voxel-wise tract-based spatial statistics (TBSS) [11] approach allows for comparisons of FA in “skeletons” within the brainstem. While this method can be fully automated, the TBSS approach analyzes data at selected high-FA voxel locations that do not necessarily belong to actual fiber tracts, and the measures at voxel level could additionally be inaccurate when FA is disrupted in the presence of disease. There are, however, increasing clinical needs for investigating brainstem fibers and their mnemonic functional relevance. A recent study [12] has established a human brainstem fiber atlas based on tractography from a large population of 488 young healthy subjects. This tractographic atlas may indeed benefit clinical neurosurgical interventions, but the neuropathological imaging applications of measuring FA in non-invasively isolated brainstem fibers are yet unknown. As atlas-based analyses are sensitive to individual uncertainties of small fibers, tractography based on native DTI data is still needed. Furthermore, the atlas was derived from data collected from healthy young adults using a high angular resolution diffusion imaging (HARDI) sequence [13], which is very difficult to implement in most clinical DTI scan protocols for neurological purposes. A reliable, clinically applicable, brainstem tractography and quantitative measure is thus needed.

The aims of this study were to 1) demonstrate the practical feasibility and neuroanatomical consistency of tractography for most trackable brainstem fibers based on a clinical DTI scan protocol; 2) automate the described tractographic procedure and test its reliability of quantitative measures of the brainstem fibers together with manual tractography; and 3) use automated tractography to assess the association between the integrity of brainstem tracts and chronic pain in an attempt to identify brainstem pathways that are specifically involved in pain regulation and processing.

## Materials and Methods

### Participants and Pain Evaluation

Seventeen participants, age between 39 and 59 years (mean age = 49.7 ± 5.2, 16 males), without brainstem lesions on MRI, were selected from patients who were recruited at the California War Related Illness and Injury Study Center (WRIISC-CA) at the Veterans Affairs Palo Alto Health Center (VAPAHCS) between March and October of 2017. All participants were combat veterans, 9 of whom were deployed to the 1991 Persian Gulf War, 6 to Afghanistan and/or Iraq as part of Operations Iraqi Freedom and Enduring Freedom, and 2 to other combat zones. All participants had endorsed diffuse combat-related symptoms via self-reported questionnaires that meet the criteria for chronic multi-symptom illness (CMI) [14]. Twelve veterans, within 24 hours before their MRI scans, had filled out item 6 from the Brief Pain Inventory Short Form (BPI) [15], which involved self-report of their current pain level (‘pain right now’). All veterans also had filled out self-reported questionnaire of the worst pain they can recall in a month prior to MRI (‘worst pain in last month’). Both pain scores were leveled from scale from 0 (none) to 10 (extreme), as an overall pain level regardless of locations. All aspects of the study were approved by the Stanford University and VAPAHCS Institutional Review Boards, and written informed consent to analyze clinical data was obtained from all participants.

### Structural MRI and DTI acquisition

Brain imaging data were collected at the Veterans Affairs Hospital in Palo Alto, CA, USA, using a 3 tesla GE Discovery MR750 scanner with an 8 channel GE head coil. High-resolution T1-weighted images (T1WI) were acquired using three-dimensional spoiled-gradient recalled acquisitions (3D-SPGR) in steady state (136 sagittal slices, TR/TE = 7.3/3.0 ms; flip angle = 11^◦^; field of view = 250 mm; slice thickness = 1.2 mm with 0.6 mm slice gap; acquisition matrix = 256 × 256; number of excitations = 1.0; resolution = 1.05 mm × 1.05 mm × 0.60 mm). T2-weighted images (T2WI) were acquired using fast spin-echo (FSE) sequences with TR/TE = 7652/98.4 ms, with 0.45 x 0.45 mm^2^ in-plane resolution and 3.5 mm slice thickness, for 47 axial slices. The T2WI images were collected for correcting geometric distortions of DTI relative to the structural T1WI. Diffusion tensor imaging scans were acquired with a 2D single-shot EPI sequence with TR/TE = 6600/84.1 ms, 1 x 1 x 2.5 mm^3^ resolutions, with 59 contiguous axial slices to enable full brain coverage. Ten scans without diffusion gradients (b0) and scans along 60 sensitization directions using diffusion-weighting gradients (DWI) with b-value of 1000s/mm^2^ were acquired for DTI reconstruction.

### DTI Processing Pipeline

The T1WI and DTI scans were initially checked for visual artifacts, and subsequently processed using an in-lab image processing pipeline, shown in Fig 1. After quality control, structural (T1WI/T2WI) and DTI data sets collected from 300 WRIISC-CA participants, who were veterans aged between 30 and 80 years, were processed through this pipeline to create customized templates and atlases in Montreal Neurological Institute (MNI) space. Creating customized templates using averaged images from the same study cohort (e.g. WRIISC-CA) is recommended [16,17], because the minimal cross-subjects variation results largely improve the registration accuracy. This in-lab pipeline included the following: 1) correction for head motion, susceptibility artifacts, and eddy-current distortions in native space using scripts from the Stanford Vision Imaging Science and Technology Lab (VISTA Lab; https://vistalab.stanford.edu/software/). Maps of DTI metrics, such as fractional anisotropy (FA), mean diffusivity, radial diffusivity, axial diffusivity, were then computed using FDT software in FSL (https://fsl.fmrib.ox.ac.uk/fsl/fslwiki/FDT). 2) Geometric distortions of DTI relative to anatomical MRI (T1WI and T2WI) in subject’s native space, which are particularly apparent in the brainstem (see supplementary S1 Fig), were corrected using Advanced Normalization Tools (ANTs) [18], through the following procedure: The initial b0 image was aligned to same subject’s T2WI using a smooth deformation algorithm and both images were subsequently transformed to anatomical T1WI using affine transformation. After distortion correction, the non-linear distortions between DTI frame and anatomical T1WI were effectively reduced. Thus, the accuracy of anatomical landmark assignment within individual spaces was improved (Supplementary S1 Fig). 3) Using nonlinear deformation algorithms in ANTs, all participants’ T1WI were diffeomorphically warped to the MNI space and averaged to create customized T1 and DTI templates. Then, an atlas with anatomical parcellations of whole-brain gray matter, white matter regions defined according to the JHU-DTI-MNI (Type I WMPM) [19] was spatially warped onto the customized template to provide anatomical labels for aiding the definition of regions-of-interest (ROIs) onto the templates. 4) The inverse of the above transformations was then used to transform anatomical parcellations and regions-of-interest (ROIs) specified in the MNI space to subjects’ native DTI space.

**Fig 1.**
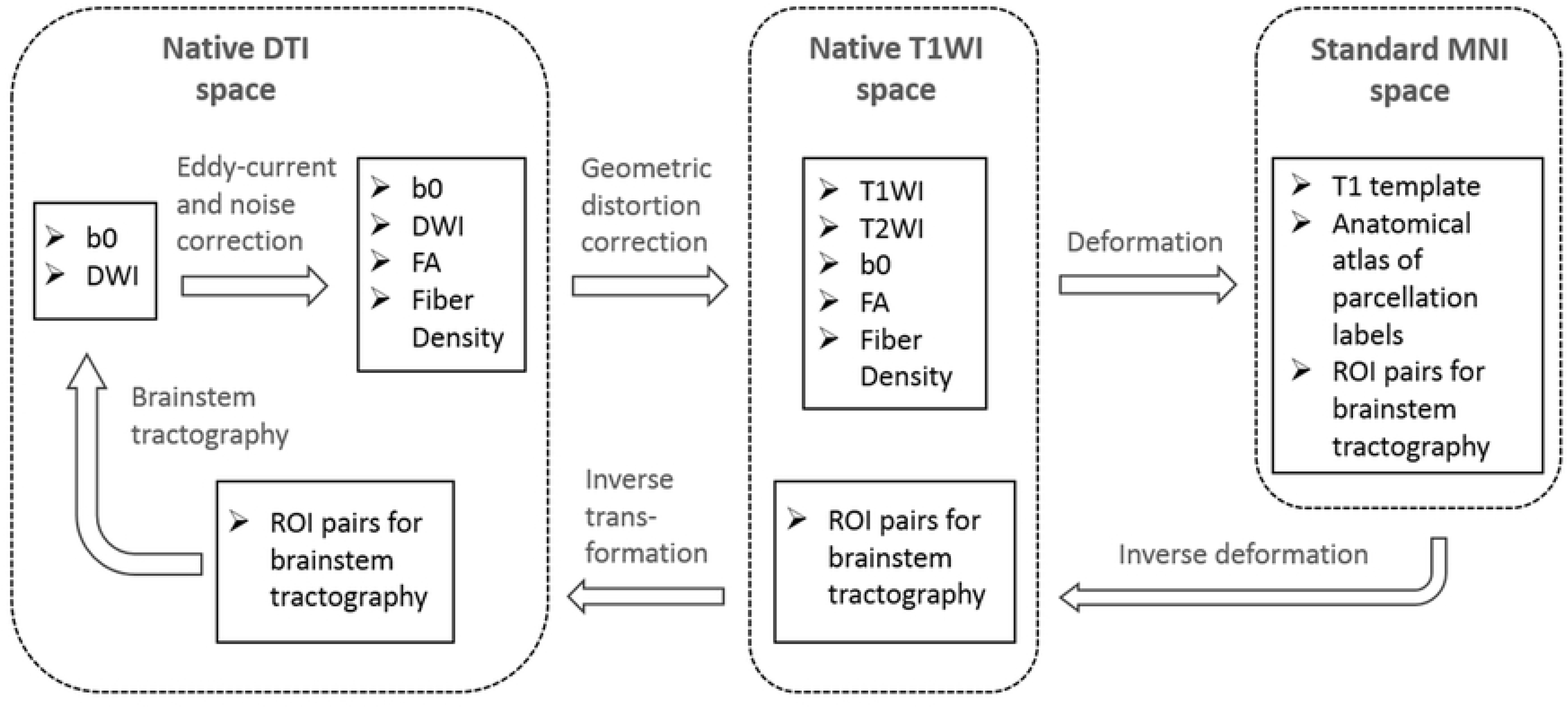
DTI processing and brainstem tractography flowchart. In-lab image processing pipeline for registration and atlas building based on structural MRI and DTI from 300 WRIISC-CA participants.

### Manual Tractography of The Brainstem Fibers

Manual brainstem tractography was performed on original b0 and DWI data using Diffusion Tensor Toolkits (DTK) (http://dti-tk.sourceforge.net/pmwiki/pmwiki.php) and TrackVis software (http://www.trackvis.org/), with a default linear least-squares fitting method in DTK. As a default function, DTK/TrackVis computed mean FA and fiber density (i.e. the streamline numbers) in each voxel of a fiber tract that resulted from tractography as the outcome measures. Manual tractography was performed using TrackVis. Each brainstem tract was initiated by placing a ‘seed’ ROI in the brainstem, and a ‘target’ ROI in the distant end. Deterministic tractography using a fixed step-length streamline propagation [20] algorithm was performed with an ‘AND’ option to reconstruct trajectories between both ROIs, with the minimum length and curvature thresholds of 10 mm and 30º, respectively. Seeds and targets were manually drawn by an experienced radiologist in accordance to known brainstem anatomy. The initial tractography consistently resulted in the intended trajectory, but additionally also resulted in other trajectories, some of which potentially being artifacts, and some that could be other neighboring pathways. In these cases, manual adjustments of ‘NOT’ ROIs were used to eliminate them. On the other hand, if the initial tractography had no trajectory results because the ‘seed’ and ‘target’ ROIs were too far apart, the distance between the ROI pairs was adjusted within their anatomical landmarks until any trajectories consistent with known anatomy could be achieved. Because manual tractography is extremely time-consuming, DTI data of 7 participants randomly selected from all participants were processed with manual tractography. Fig 2 depicts the ROI placements and reconstructed streamlines of nine fiber tracts that travel through the brainstem. For comparison with sectional anatomy, cross-sectional slices at the lower and upper brainstem panel of the color-oriented anisotropic map, six representative brainstem fibers, and corresponding neuroanatomy are shown in Fig 3. Overall, nine pairs of brainstem fiber tracts were tracked:

**Fig 2.**
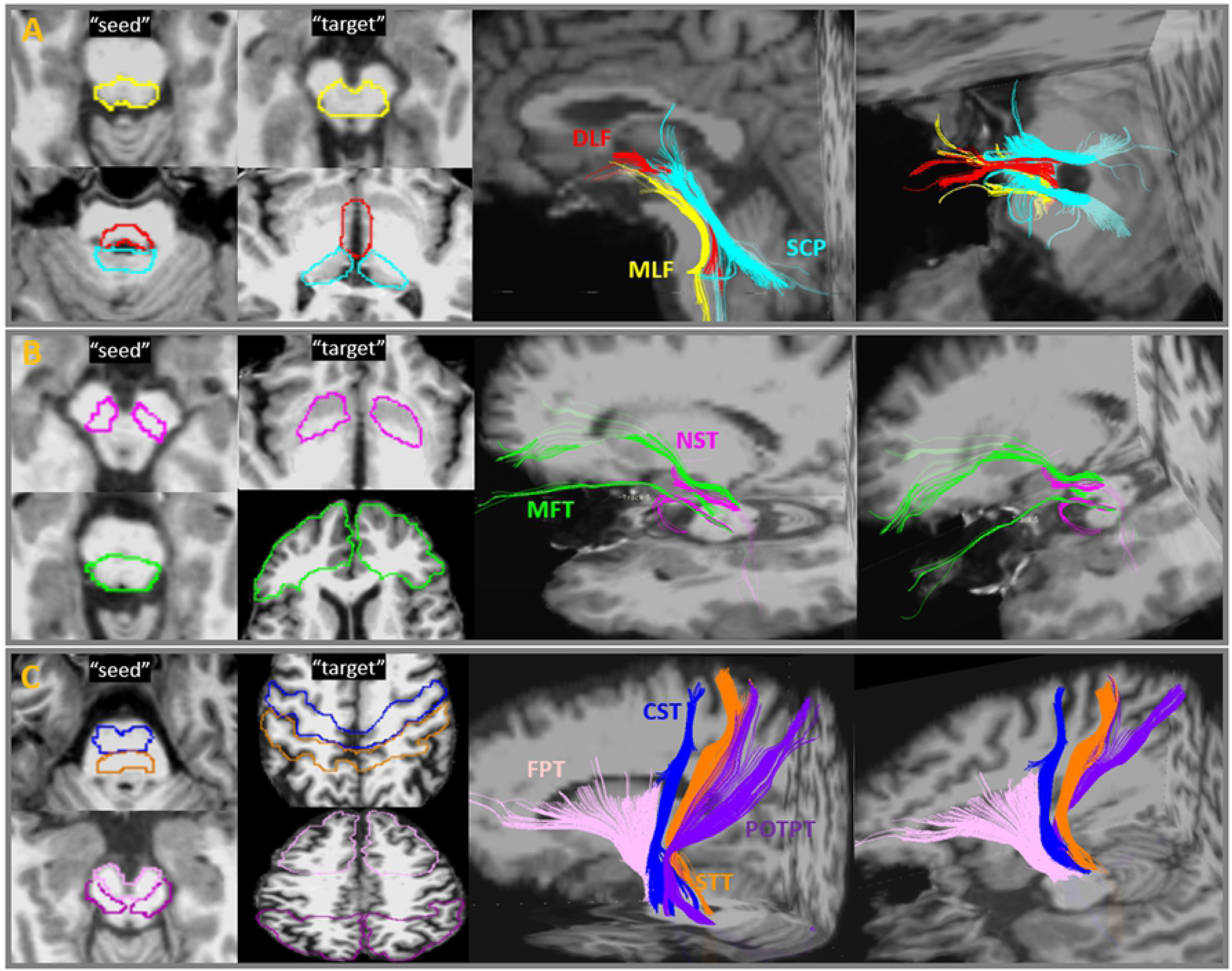
Brainstem fiber tracts and manual ROI placement. Placement of ROI-pairs (“Seed” and “target”) and their fiber outputs performed on TrackVis. Abbreviations: MLF= medial longitudinal fasciculus; DLF = dorsal longitudinal fasciculus; SCP = superior cerebellar peduncle; MFT = medial forebrain tract; NST = nigrostriatal tract; FPT = frontopontine tract; CST= corticospinal tract; STT = spinothalamic tract; POTPT = parieto-, occipito-, and temporopontine Tract.

**Fig 3.**
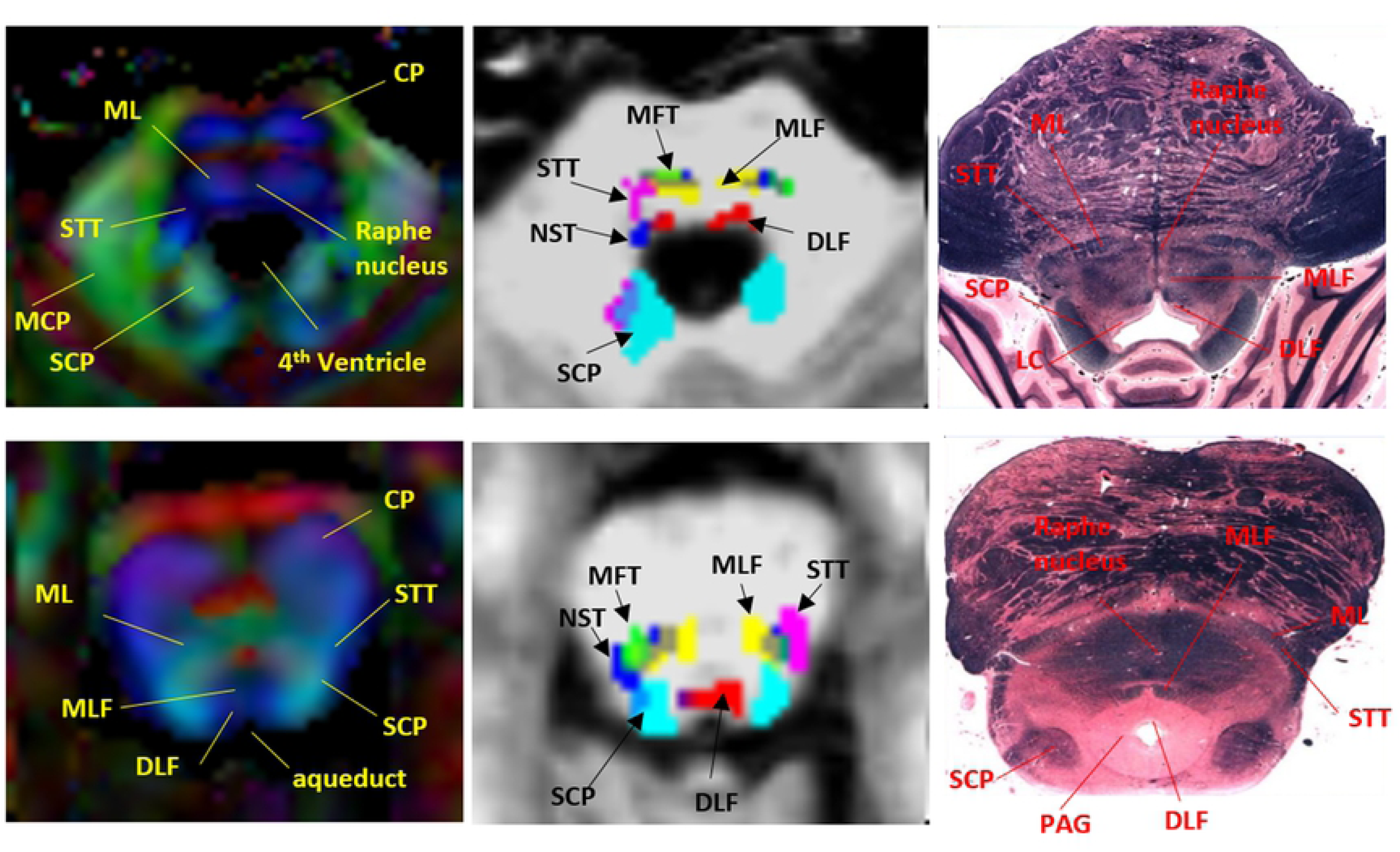
Representative brainstem fiber tracts and corresponding neuroanatomy in two brainstem levels. *Left*: color-coded anisotropic map (red: left-right oriented; blue: superior-inferior oriented; green: anterior-posterior oriented); *Middle*: six brainstem reticular tracts superimposed on a T1WI; *Right*: corresponding anatomic sections. Abbreviations: CP = cerebral peduncle; ML= medial lemniscus; STT = spinothalamic tract; MCP = middle cerebellar peduncle; SCP = superior cerebellar peduncle; MLF= medial longitudinal fasciculus); DLF = dorsal longitudinal fasciculus; MFT = Medial Forebrain Tract; NST = nigrostriatal tract; LC =locus coeruleus; PAG = periaqueductal gray matter. The brainstem anatomical pictures are downloaded from the web site (http://www.dartmouth.edu/~rswenson/Atlas/BrainStem/index.html) with permission of the author.

1. Medial longitudinal fasciculus (MLF; Fig 2A. **–** yellow): the seed ROI was placed in the area anterior to the 4th ventricle floor, including the inferior cerebellar peduncle (ICP) and the medial lemniscus (ML). The target ROI was placed in the midbrain (MidB), encompassing the ventral tegmentum area (VTA) and the red nuclei (RedN). In the brainstem, the reconstructed fibers travelled ventrally to the aqueduct and the 4th ventricle, slightly laterally to the midline, where the Raphe nuclei are located (Fig 3). This pattern was consistent with previous tractography of healthy human brainstems and known anatomy of MLF [12,21].
2. Dorsal Longitudinal Fasciculus (DLF; Fig 2A – red): the seed ROI was placed in the area anterior to the aqueduct, including the periaqueductal gray matter (PAG) and surrounding area as well as the locus coeruleus (LC). The target ROI was placed in the dorsal MidB, including the mammillothalamic tract, which is located laterally to the cerebral aqueduct. The reconstructed fibers connected between the LC and the hypothalamus, travelled ventrolaterally to the cerebral aqueduct, and ran dorsocaudally to the MLF (Fig 3). The tractography pattern of DLF was consistent with known anatomy and a previously-described tractography atlas [12,21].
3. Superior Cerebellar Peduncle (SCP; Fig 2A – cyan): The seed ROI was placed in the area posterior to the 4th ventricle floor and aqueduct, including white matter in the superior cerebellar peduncle and the cerebellar dentate. The target ROI was placed in the ventral thalamus. The reconstructed fibers connected the dentate nuclei and the thalamus, including non-decussated and decussated dentato-rubrothalamic tracts. This tract and its anatomy has been reported in many previous tractography studies of the human brain [12, 22].
4. Nigrostriatal tract (NST; Fig 2B – pink): The seed ROI was placed in the MidB, including the entire substantia nigra (SN). The target ROI was placed in the inferior level of the putamen and the globus pallidus formation (Put/GP). This small fiber travelled from the SN dorsomedially to the subthalamic nuclei (STN) and turned dorsolaterally toward the ventral Put/GP. This pattern was consistent with previous tractography findings and known anatomy of the NST [23].
5. Medial Forebrain Tract (MFT; Fig 2B – green): The seed ROI was placed in the ventral MidB area anterior to the aqueduct, including ventral tegmental area (VTA). The target ROI was placed in the inferior frontal gray and white matter area, including the rectus, orbitofrontal, and the prefrontal cortices. This tract originated in the VTA, extended dorsolaterally toward the nucleus accumbens (NAc) and anterior thalamic radiation (ATR, located in the anterior limb of the internal capsule), and finally projected to the orbitofrontal cortex (OFC) and the dorsolateral prefrontal cortex (DLPFC). The tractography and neuroanatomy of MFT has been demonstrated in previous papers [24,25].
6. Frontopontine Tract (FPT; Fig 2C – light pink): The seed ROI was placed in the ventral medial 1/3 of the cerebral peduncle (CP). The target ROI was placed in the middle and superior frontal gray and white matter, including the supplementary motor and the premotor areas, but excluding the primary motor area. The FPTs, originate from the frontal lobe, descend through the anterior limb of the internal capsule, and end in the pontine nuclei, have been described in many neuroanatomy textbooks and tractography studies [26].
7. Corticospinal Tract (CST; Fig 2C – blue): The seed ROI was placed in the middle 1/3 of the CP. The target’ ROI was placed in the precentral gray and white matter, which corresponds to the primary motor area. The CST is the primary motor pathway descending from the motor cortex innervating muscles controlling skilled voluntary movement. This ROI placement of tracking the CST is in line with previously well-described strategies to reconstruct CST using DTI tractography [26].
8. Spinothalamic Tract (STT; Fig 2C – orange): The seed ROI was placed in the ML and the surrounding area. The target ROI was placed in postcentral (sensory) gray and white matter, corresponding to the primary somatosensory area. The STT rose up the brainstem ML area, projected to thalamus, and eventually reached the sensory cortex. The tractography of this pathway has been demonstrated previously [12].
9. Parieto-, Occipito-, and Temporopontine Tract (POTPT; Fig 2C – purple): The seed ROI was placed in the laterocaudal third of the CP. The target ROI was placed broadly in the parietal, occipital, and temporal gray and white matter parcellations. The parietopontine and occipitopontine tracts passed the posterior limb of the internal capsule, retrolenticular part of the internal capsule (RLIC), and the dorsal thalamus; the temporopontine tract travelled medially to the hippocampoamygdala formation. These patterns were in agreement with a previous report [12].

The tractographic anatomy, possibly related neurotransmitter systems and functions of the nine tracts reconstructed from diffusion tensor tractography are summarized in Table 1.

**Table 1.** Anatomy, possible associated neurotransmitters and functions of brainstem circuits.

| Brainstem tracts | Gray/white matter structures in the brainstem tractographic pathways | Possible Neuro-transmitter | Related Functions |
| --- | --- | --- | --- |
| MLF | Begin: ICP and ML;<br>Pass: lower raphe nuclei, ML;<br>End: MidB and RN | Serotonin,<br>Noradrenaline | wakefulness, pain<br>control, maintaining<br>posture, cardiovascular<br>control, eye movement |
| DLF | Begin: PAG and Surrounding area;<br>Pass: LC, PAG;<br>End: hypothalamus; mamillary body | Serotonin,<br>Noradrenaline,<br>Histamine | wakefulness, attention,<br>consciousness, stress and<br>reward, pain |
| SCP | Begin: SCP and cerebellar dentate nuclei;<br>Pass: PPN;<br>End: RN and thalamus | Acetylcholine,<br>GABA | sleep, cognition, mood,<br>attention, arousal,<br>voluntary limb<br>movements, locomotion |
| NST | Begin: SN<br>Pass: VTA, STN<br>End: putamen and GP | Dopamine,<br>Serotonin | motor function, mood,<br>pleasure and<br>reinforcement |
| MFT | Begin: MidB and VTA<br>Pass: NAc, ATR<br>End: rectal, orbitofrontal, and<br>prefrontal cortices | Acetylcholine | mood, word, reward and<br>pleasure |
| FPT | Begin: medial 1/3 of the CP<br>Pass: ALIC, SCR, frontal cortices<br>End: superior frontal, supplementary motor<br>area, and the premotor areas | Non-reticular<br>system | control of movement,<br>facial movement |
| CST | Begin: middle 1/3 of the CP<br>Pass: PLIC<br>End: precentral (primary motor) cortex | Non-reticular<br>system | movement of body<br>muscles |
| STT | Begin: ML<br>Pass: dorsal spinal cord, ML, thalamus<br>End: postcentral (sensory) cortex | Non-reticular<br>system | sensory of temperature,<br>pain |
| POTPT | Begin: lateral 1/3 of the CP<br>Pass: ventral medulla, RLIC, dorsal thalamus,<br>amygdala and hippocampus<br>End: parietal, occipital, and temporal cortex | Non-reticular<br>system | variety of functions:<br>working memory,<br>learning, visual, auditory |

Abbreviations: ICP = inferior cerebellar peduncle; ML = medial lemniscus; MidB = midbrain; RN = red nucleus; PAG = periaqueductal grey matter; LC = locus coeruleus; SCP = superior cerebellar peduncle; PPN = pedunculopontine nucleus; SN = substantia nigra; VTA = ventral tegmental area; STN = subthalamic nucleus; GP = Globus pallidus; NAc = Nucleus Accumbens;
ATR = anterior thalamic radiation; CP = cerebral peduncle; ALIC = anterior limb and genu of the internal capsule; SCR = superior coronal radiata; PLIC = posterior limb of the internal capsule; RLIC = retrolenticular part of the internal capsule.

### Automated Brainstem Tractography

A common practice of automated-tractography is to apply pre-defined ROI-pairs from the MNI atlas space onto individual DTI data to extract only those streamlines that run through both ROI ends. However, most of the brainstem fiber tracts (e.g. MLF, DLF, NST and MFT) do not have a previously established ROI atlas or parcellations in MNI space. In this study, we use the manually-defined ROI-pairs and fiber results to build the pre-defined ROI templates of the brainstem tracts and subsequently tested the reproducibility of these tractography results in comparison with manual tractography. Briefly, the procedure of building the templates of ROI-pairs in MNI space included: 1) employing the in-lab imaging processing pipeline to transfer the manually defined ROI pairs and the 9-brainstem fibers into MNI space. 2) in MNI space, the normalized ROI pairs of each fiber were averaged and binarized to create the pre-defined ROI-pair templates; the normalized fibers were also averaged to create fiber templates as the anatomical references; 3) to further improve anatomical precision of each fiber isolation, minimal adjustments of the ROI templates were performed including: (i) dilating the ROI boundaries to ensure the ROI-pairs fully cover the termini of each fiber template, (ii) eroding ROI boundaries between two adjacent ROIs (e.g. the boundaries of MLF and DLF) to eliminate overlapping tractographic results; (ii) limiting the dilation and erosion of the ROI templates within the anatomical landmarks based on the JHU-DTI-MNI atlas (also described in Table 1 as the ‘begin’ and ‘end’ regions). Once the templates of ROI-pairs were built in MNI space, they were transferred backwards through the inverse transformation of the in-lab image processing pipeline onto all participants’ native DTI space to launch individual brainstem trajectories using TrackVis between the assigned ROI-pairs. The overall processing time of this automated tractography procedure for tracking the 9 brainstem fiber pairs was approximately 30~40 minutes per subject.

### Test-retest of the Quality and Quantity for the Brainstem Tracts

There is currently no comparative standard for the assessment of anatomical precision of tractographic approaches. Defining false positive or negative fiber tracts from tractographic results relies on prior anatomical knowledge. Therefore, tractography with ROIs that have been manually defined by experts using such anatomical knowledge and extensive training, although imperfect, is a conventionally accepted validation method.

While the presence of unexpected fiber lines (i.e. false positive fibers) can be removed by the tractographer based on prior anatomical knowledge during tractography, the absence of expected fiber lines (i.e. false negative fibers) is still a problem that can result in failure of quantitative measurements along existing fiber lines. In this study, we tested the ‘success rate’ for each brainstem fiber in terms of the subjects’ numbers that successfully presented a certain brainstem fiber, divided by the subjects’ numbers that brainstem tractography was performed based on either manual or automated tractography. The ‘success rate’ reflects the feasibility of brainstem tractography in its utilization for quantitative measurement.

Next, two quantitative scalar indices were obtained in each presented brainstem tract as the major outcome measures: (1) The FA index, which indicates the microstructural integrity of fiber connection, based on the mean FA per 1 mm^3^ voxel in each brainstem tract. The fiber density index, which represents the weights of reconstructed fibers, based on the mean streamline numbers per 1 mm^3^ voxel in each brainstem tract [27]. We used an intra-class correlation coefficient (ICC) to test if quantitative measures based on automated tractography could reliably reproduce the same measure based on manual tractography. The reproducibility explains how reliable certain outcome measures based on automated tractography could be applied as an alternative approach to manual tractography.

## Results

### Success Rate and Reproducibility of Quantitative Tractography

Supplementary Table S1 lists the success rates of manual and automated tractographic performances in each brainstem tract. For manual tractography, most of the brainstem fibers were successfully detected. Some failed tracking occurred in fibers that travelled through long distances or areas susceptible to artifacts, crossing-fibers and other quality problems. Automated tractography achieved a slightly smaller, but similar success rate as the manual tractography. Both approaches had 85-100% success rates, which allow for the outcome measures and analyses based on our conventional DTI quality.

Supplementary Table S1 also summarizes reproducibility between manual and automated tractographic approaches. The FA in fibers measured by automated tractography reliably reproduced those measured by manual tractography. On the other hand, fiber density from automated tractography could marginally reproduce the results from manual tractography.

### Relations between DTI and Pain levels

Linear regression models were used to examine the relationships of FA or fiber density in each brainstem tract with the two self-reported pain levels (i.e. ‘pain right now’ and ‘worst pain last month’), separately. Table 2 and Fig 4 show that, significant associations were observed between both pain levels and FA in the DLF and MLF, but not in other brainstem tracts. Specifically, after controlling for age and gender, per unit increase in ‘pain right now’ level corresponded to a 3.48% (CI_95_: 0.5 to 6.4%, *p* = 0.026) FA reduction in DLF and a 3.02% (CI_95_: 0.7 to 5.4%, *p* = 0.017) FA reduction in MLF. Furthermore, per unit increase in ‘worst pain last month’ corresponded to a 2.30% (CI_95_: 0.7 to 3.9%, *p* = 0.008) FA reduction in DLF and 1.87% (CI_95_: 0.4 to 3.4%, *p* = 0.019) FA reduction in MLF. No significant relation between pain levels and fiber density was detected.

**Table 2.** Linear regressions between FA of brainstem fibers and pain levels.

| Tract<br>name | Pain Right Now<br>(n=12) |  |  | Worst Pain Last Month<br>(n=17) |  |  |
| --- | --- | --- | --- | --- | --- | --- |
|  | Mean (S.E.) % |  |  | Mean (S.E.) % |  |  |
| | FA change rate<br>/scale increase | Effect-size | $p$-value | FA change rate<br>/scale increase | Effect-size | $p$-value |
| **MLF** | **-3.02 (1.04)** | **-1.75** | **0.017** | **-1.87 (0.71)** | **-1.32** | **0.019** |
| **DLF** | **-3.48 (1.30)** | **-1.61** | **0.026** | **-2.30 (0.75)** | **-1.53** | **0.008** |
| **SCP** | -7.32 (3.28) | -1.34 | 0.053 | -2.98 (1.47) | -1.02 | 0.062 |
| **NST** | -0.72 (1.50) | -0.15 | 0.644 | -2.47 (1.83) | -0.75 | 0.204 |
| **MFT** | 0.94 (0.62) | 1.07 | 0.180 | 0.30 (0.53) | 0.33 | 0.582 |
| **FPT** | -0.60 (0.82) | -0.47 | 0.482 | 0.35 (0.63) | 0.29 | 0.588 |
| **CST** | 0.21 (1.05) | 0.12 | 0.860 | 0.84 (0.95) | 0.51 | 0.393 |
| **STT** | -0.59 (0.66) | -0.53 | 0.400 | 0.01 (0.68) | 0.01 | 0.989 |
| **POTPT** | -0.79 (1.39) | -0.38 | 0.586 | -0.04 (1.12) | -0.02 | 0.970 |

Mean (S.E.) FA change rate (%) associated with per increased pain scale, effect size and significance that estimated based on linear regression test for each brainstem fiber.
Effect size = $(2 * T\text{-value}) / \text{SQRT}(\text{degree of freedom})$

**Fig 4.**
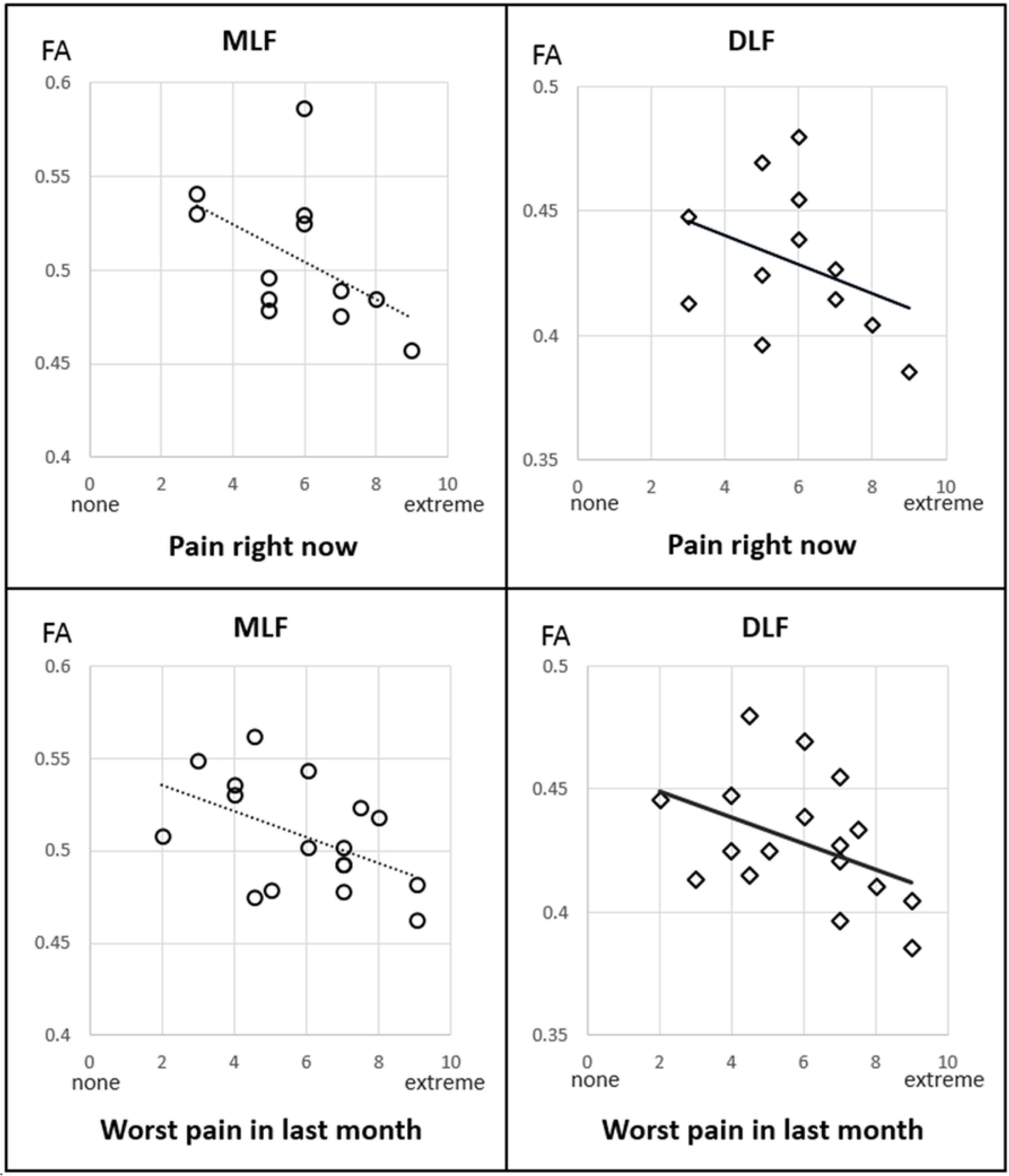
Relations between FA and Pain levels. Scatter plots of relations between FA of the MLF and DLF tracts and two pain scales.

## Discussion

In this study, we introduced an automated tractographic approach based on a predefined ROI atlas for efficiently tracking and isolating small white matter tracts in the human brainstem. Our results demonstrated the usefulness of quantitative tractography in analyzing the relationship between brainstem tracts and chronic pain. This application was used to detect several brainstem fibers and diminished WM integrity levels were associated with pain severity.

### Brainstem Tractographic Anatomy and Neurological Implications

Most brainstem fibers could be subdivided into three types of circuits: 1) the corticopontine fibers that travel between spinal cord, pons, and the neocortices, including FPT, CST, STT, POTPT (Table 1 and Fig 2C); 2) the reticular tracts that interconnect nuclei that are located throughout the brainstem and the midbrain, including DLF, MLF, SCP, MFT and NST (Table 1 and Fig 2A and 2B); and 3) the pontinopeduncle tracts that run transversely between pons and cerebellar peduncles including the middle cerebellar peduncle, the inferior cerebellar peduncle and pontine crossing tracts. Among this traffic, the reticular tracts have major clinical significance in indicating essential neurotransmitter functions (Table 1 and Fig 5). Manual and automatic tractography of the corticopontine fibers and pontinopeduncle tracts have been well-established previously. By contrast, tractography of the reticular tracts needs to be further established as these fibers are thought to contain a variety of autonomic functions through corresponding neurotransmitter pathways.

**Fig 5.**
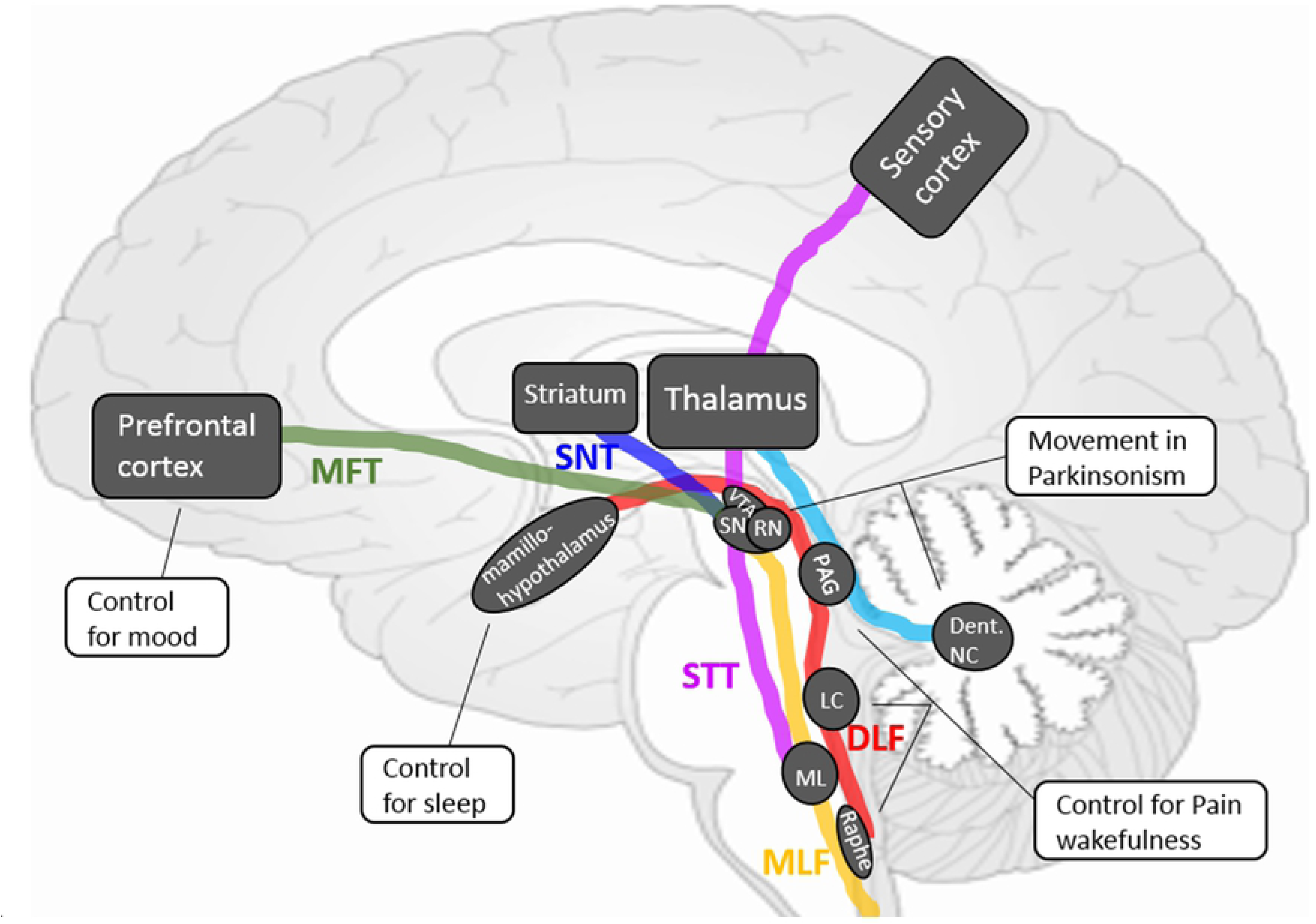
Illustration of brainstem reticular tracts. Conceptual sketch according to the DTI tractography of the brainstem reticular tracts, their anatomy, fiber-linked gray matter centers and potentially associated functions.

MLF is generally known to connect the cranial nerves (III, IV, and VI) which are involved in eye movement, as well as the stabilization of the head and neck. MFL is also known to send parasagittal fibers from the dorsal raphe nuclei, which serve as a center of serotonergic neurons [28]. The role of MLF in modulating some autonomic, reproductive and excretory functions, as well as modulating perception to pain needs to be investigated. The DLF that links the hypothalamus, periaqueductal gray matter, locus coeruleus, and the tractus solitarius in the medulla, carries both ascending and descending fibers. This tract is considered to be a pathway regulating the sleep–wake cycle, mood, and cognition [29,30]. DLF is also related to pain control [31]. SCP is a large bundle of projection fibers arising chiefly from the cerebellar dentate nuclei, ascending along the posterior floor of the forth ventricle, and ending in the red nucleus and thalamus. SCP plays a major role in the coordination of movement of the ipsilateral limbs. Previous studies [32,33] also suggest that SCP contains GABAergic fibers. The NST is a dopaminergic pathway that is involved in Parkinson’s disease. Individual tractography-guided quantitative DTI measures of this tract have been suggested as a promising marker for identifying Parkinson’s disease and such measurements may correlate with the severity of motor dysfunction [34–36]. NST is also suggested to control depressive fatigue states in Parkinson’s Disease [37,38]. MFT, which connects the brainstem to nucleus accumbens, anterior thalamic radiation, and dorsal and medial prefrontal cortices, is involved in mood, the reward system, the appetitive motivational seeking system and euphoria [39]. The relation of MFT and major depressive disorder has been well-documented through DTI [24], and MFT tractography is also used for targeting deep brain stimulation in depression [25].

Given the variety of the reticular fibers and their corresponding functions, brainstem DTI tractography may be highly informative in the diagnosis and study of a range of neurological conditions. Our newly developed brainstem tractographic approach allows the correlation analyses for clinical measures and quantitative DTI measures of fiber tracts known to be involved in those neurological dysfunctions.

### Quantitative Brainstem Tractography and Pain

Much of the DTI and FA studies have been focusing on examining differences in white matter microstructure in chronic pain vs. non-pain controls, and correlating FA with pain levels (see a recent review paper [40]). Based on TBSS, pain-related FA reductions have been detected in multiple white matter tracts along diffuse ‘white matter skeletons’, including the internal and external/extreme capsules, corpus callosum, cingulum, thalamic radiation, and brainstem white matter, primary somatosensory and motor cortices, orbitofrontal cortex [41,42], rostral anterior cingulate, and dorsal lateral prefrontal cortex [43]. TBSS and most voxel-based analytic approaches, however, are known to be highly sensitive to registration errors and may produce false positive findings [44]. Moreover, alterations in most cerebral white matter regions could also be observed in comorbid neurological symptoms such as cognitive or mood impairments, but are not specifically responsible for pain [45]. In this context, it is essential to identify key white matter circuits involved in pain perception and regulation, and to explore the potential disruptions of these key pathways in relation with pain severity levels. Electrophysiological [46] and functional MRI studies [47] have shown that stimulus-evoked pain is associated with activity in several brain areas (‘pain centers’) and pathways linking these areas. However, these observations did not address the actual tracts and their structural connections in association with pain states. According to brainstem anatomy, both the ‘ascending (bottom-up) pathway’ – which sends the nociceptive flow from the spinal cord to cerebral cortices, and the ‘descending (top-down) pathway’ – that supports a top-down role in modulating the nociceptive signals, are involved in the brainstem fiber traffic [48]. A previous DTI study [49] demonstrated that fiber tracts originating in the periaquaductal grey (PAG) that project towards other brain areas, which may represent the descending pain control pathway, can be successfully visualized using probabilistic tractography. however, this study did not provide specific fiber isolation and quantitative measures such as FA in subjects with pain. Our study further isolated this theoretical ‘descending pathway’ to mainly overlap with the DLF and MLF tracts. We found that diminished FA in these tracts is associated with both ongoing pain (i.e. ‘pain right now’) and chronic pain (i.e. ‘pain in last month’) levels. These findings, suggest that quantitative brainstem tractography is a promising marker for investigating pain regulation mechanisms. Further analyses based on larger patient populations and healthy controls are needed to validate these preliminary findings.

### Advantages of Automated Brainstem Tractography

This newly developed automated brainstem tractography has several merits: First, this approach has been shown feasible to be applied, implemented and performed on conventional DTI scans. Tracking the brainstem functional fibers on DTI is known to be difficult, and the detection of those small fiber anatomies depends on to the quality of acquisition including resolution, directions, controlling of noise and crossing-fibers. Existing approaches that provide comprehensive brainstem tractography have been established based on sophisticated diffusion acquisitions such as HARDI, diffusion spectrum (DSI) [50], etc., which provide limited clinical utility. In this study, we applied brainstem tractography using a readily available and sufficiently short (8 min) conventional DTI sequence, and demonstrated efficient success rates based on either manual or automated tractographic approaches. This suggest that the automated tractography could be adopted for quantitative measures in most clinical DTI protocols that use conventional resolutions (4~8 mm^3^ voxel size) and directions (30~80). Second, this automated tractographic approach provides reproducible measures that overcome known difficulties associated with manual tractography. Manual brainstem tractography is time consuming, depends on individual experience, and may suffer from reproducibility problems. Most existing automated tractography programs [10,51–53] have been used for representing tract trajectories of several major cerebral pathways with known anatomy. Automated brainstem tractography is currently not available due to a lack of atlas-segmented ROIs of brainstem small regions. Our study developed this automated tractographic approach based on pre-defining ROI pairs in the standard MNI space, with a combination of a high-quality DTI and MRI image registration pipeline. We demonstrated that this approach achieves similar FA measures as manual tractography while offering significant benefits through elimination of subjective rater bias, greatly reduced processing times, and the feasibility for quantitative analyses of large numbers of DTI data.

## Limitations

The work presented in this paper has several limitations: first, we did not identify all brainstem tracts: 1) Tractographic visibility is subject to the quality of DTI scans. Small tracts, such as the rubrospinal and the central tegmental tracts, previously addressed by Meola and colleagues [12], are not identifiable due to the limited quality of the DTI scan protocol used herein. 2) Other pain-specific fibers connecting the PAG to cerebral cortices, described by Hadjipavlou and colleagues [49], were not included in our approach, because the anatomical certainty and practical feasibility of these fibers need to be validated yet. 3) We did not include the pontinocerebellar, pontinomedulla, pontinospinal tracts, or cranial nerves into this approach due to the coverage of the brain scans and consideration of the susceptibility artifacts near the skull-base that may confound local fiber tracking. 4) We also did not perform tractography of the middle or the inferior cerebellar peduncles, as well as the pontine crossing tracts because of a lack of knowledge regarding the physiological and functional relevance of these transverse fibers. Another limitation is that we did not test other microstructural measures (e.g. mean, radial, axial diffusivities) in the brainstem fibers, although these measures might provide complementary information about fiber integrities. Nevertheless, quantitative DTI changes of specific brainstem fibers provide a promising biomarker in chronic pain. Further validation is warranted using this automated tractographic approach in larger populations, including more detailed pain evaluations and a more comprehensive consideration of all available DTI metrics.

## Conclusions

This study developed an automated approach for tracking brainstem fibers based on DTI. This approach is feasible for quantitative analyses and is an alternative to the very time consuming manual tractography. We demonstrated the utility of this approach in correlating the disruptions of some key brainstem tracts with individual pain severities. Further applications of this brainstem tractographic approach may aid in the exploration of clinical implications of these fibers in association with chronic pain syndromes, sleep disturbances, mood alterations, autonomic disturbances and other neurological disorders.

## Acknowledgement

We thank Ms. Stacy Moeder for administrating the California War Related Illness and Injury Study Center (WRIISC-CA) research programs.

## Supporting Information

**S1 Fig**. **Example of distortion corrections through image processing pipeline.** Intrinsic distortion correction between individual FA map (green color) and structural T1WI (gray color) improves registration accuracy in brainstem area (arrowed and zoomed areas).

**S1 Table. Results of test-retest for the brainstem tractographic approaches.** The success rates of each manual and automated tractographic performance, and the reproducibility of each outcome measure (FA and fiber density) between manual and automated tractography.

